# OsNLP3/4-OsRFL module regulates nitrogen-promoted panicle architecture in rice

**DOI:** 10.1101/2023.02.28.530553

**Authors:** Jie Wu, Liang-Qi Sun, Zi-Sheng Zhang, Ying Song, Yu Bai, Guang-Yu Wan, Jing-Xian Wang, Jin-Qiu Xia, Zheng-Yi Zhang, Cheng-Bin Xiang

**Author notes:** Corresponding author: Cheng-Bin Xiang). These authors contributed equally (Jie Wu, Liang-Qi Sun, Zi-Sheng Zhang).

## Abstract

Rice panicles, a major component of yield, are regulated by phytohormones and nutrients. How mineral nutrients promote panicle architecture remains largely unknown. Here, we report that NIN-LIKE PROTEIN3 and 4 (OsNLP3/4) are crucial positive regulators of rice panicle architecture in response to nitrogen (N). Loss-of-function mutants of either *OsNLP3* or *OsNLP4* produced smaller panicles with reduced primary and secondary branches and fewer grains compared with wild type, whereas their overexpression plants showed the opposite phenotypes. Notably, the OsNLP3/4-regulated panicle architecture was positively correlated with N availability. OsNLP3/4 directly bind to the promoter of *OsRFL* and activate its expression to promote inflorescence meristem development. Furthermore, OsRFL activates *OsMOC1* expression by binding to its promoter. Our findings reveal the novel N-responsive OsNLP3/4-OsRFL-OsMOC1 module that integrates N availability to regulate panicle architecture, shedding light on how nutrient signals regulate panicle architecture and providing candidate targets for the improvement of crop yield.

## Introduction

Rice (*Oryza sativa* L.), one of the most important cereal crops in the world, produces staple food for more than half of the global population (Zuo and Li, 2014). To feed the increasing global population, it is urgent and challenging to increase the rice yield. Grain number per panicle is a major contributor to rice yield, which is mainly determined by panicle architecture composed of rachis length and the arrangement of branches and spikelets (Xing and Zhang, 2010; Zhang and Yuan, 2014). Thus, it is particularly important to identify the regulatory mechanism of panicle architecture for developing larger-panicle rice varieties to improve grain yield.

The development of rice panicles initiates from the transition of shoot apical meristem (SAM) to inflorescence meristem (IM), followed by IM producing axillary meristem (AM), then differentiates into primary branch (PB) and secondary branch (SB) primordia, and finally spikelet meristem (SM), which directly determines the panicle architecture (Ikeda et al., 2004; Zhang and Yuan, 2014). During past decades, many genes that regulate panicle development have been identified. *Grain Number1a* (*Gn1a*), which encodes a cytokinin oxidase, is the first major quantitative trait locus found to be associated with panicle development. Reduced *Gn1a* expression leads to increased cytokinin content in the IM, which subsequently increases the number of branches and grains per panicle (Ashikari et al., 2005). *LAX PANICLE1* (*LAX1*), *LAX PANICLE2* (*LAX2*), *MONOCULM1* (*MOC1*) and *MOC3* act as key regulators in AM formation and maintenance. Loss of function of these genes results in partial or even complete disappearance of AM, leading to severe reduction of lateral spikelet on branches (Komatsu et al., 2003; Lu et al., 2015; Tabuchi et al., 2011; Zhang et al., 2020). In addition, ABERRANT PANICLE ORGANIZATION1 (APO1) and RICE FLORICAULA-LEAFY HOMOLOG (RFL) interact with each other to regulate the temporal fate of IM, thus increasing the number of primary branches and grains per panicle (Ikeda-Kawakatsu et al., 2012; Ikeda et al., 2007; Ikeda et al., 2005; Rao et al., 2008). Thus, the stability of the APO1-RFL interaction, which is reduced by the E3 ubiquitin ligase encoded by the *LARGE2* gene, affects panicle architecture (Huang et al., 2021). In addition, genes such as *DENSE AND ERECT PANICLE1* (*DEP1*), *SQUAMOSA PROMOTER BINDING PROTEIN-LIKE14* (*SPL14*), and *ORYZA SATIVA HOMEOBOX1* (*OSH1*) can also change the panicle architecture by affecting the length of the rachis and branch (Duan et al., 2019; Huang et al., 2009; Lu et al., 2013).

Nitrogen (N) is crucial for crop yield in part by affecting plant architecture, such as tiller number and panicle architecture (Luo et al., 2020). The application of N fertilizer before panicle initiation is known to increase the grain number per panicle (Ding et al., 2014). Moreover, grain number is linearly correlated with total plant N content, and rice genotypes with high N utilization capacity tend to produce more grains per panicle (Makino, 2011; Yoshida et al., 2006). In addition, plants overexpressing N starvation-inducible *C-TERMINALLY ENCODED PEPTIDE6.1* (*OsCEP6.1*) exhibited shorter panicles and fewer grains than the control (Sui et al., 2016). Putative nitrate-peptide transporter family genes (NPFs), such as *OsNPF7.7* and *OsNPF7.1a*, promote the assimilation of nitrate and ammonium and enhance panicle size and branches (Huang et al., 2018; Huang et al., 2019). Nevertheless, the molecular mechanisms by which N promotes panicle architecture remain largely unknown.

NIN-LIKE PROTEINs (NLPs), a subfamily of the plant-specific RWP-RK transcription factor family, play crucial roles in various aspects of plant growth and development in response to N signaling (Konishi and Yanagisawa, 2011; Mu and Luo, 2019). For instance, *AtNLP7* in *Arabidopsis* acts as a nitrate sensor to regulate the expression of multiple N-related genes through a specific nuclear retention mechanism that is regulated by phosphorylation of calcium-dependent protein kinases (CPKs) (Liu et al., 2022; Liu et al., 2017; Marchive et al., 2013; Yu et al., 2016).

*AtNLP8* is a master regulator of nitrate-promoted seed germination (Yan et al., 2016). In *M. truncatula*, MtNLP2 and MtNIN directly activate the expression of the leghemoglobin gene, thus balancing the oxygen content and promoting efficient N fixation in nodules. In addition, nitrate-mediated formation of NIN-NLP heterodimers inhibits nodulation (Jiang et al., 2021; Lin et al., 2018; Nishida et al., 2021). In maize and barely, *ZmNLP5*, *ZmNLP6*, *ZmNLP8*, *ZmNLP3.1* and *HvNLP2* also play critical roles in controlling nitrate assimilation and signaling (Cao et al., 2017; Gao et al., 2021; Ge et al., 2020; Wang et al., 2018). However, NLPs have never been reported to regulate panicle architecture.

Our previous work showed that OsNLP3 and OsNLP4 (OsNLP3/4) are key positive regulators of grain yield and nitrogen use efficiency (NUE) in rice (Wu et al., 2021; Zhang et al., 2022). We continue to explore novel functions of OsNLPs and report here that OsNLP3/4 improve grain yield by upregulating the common target gene *OsRFL* and thereby improving the panicle architecture with increased panicle length, number of primary and secondary branches, and grain number per panicle.

Importantly, OsNLP3/4-improved panicle architecture is positively correlated with N concentrations. Moreover, we identified *OsMOC1* as a novel target gene of OsRFL. OsRFL upregulates *OsMOC1* by directly binding to the promoter to control panicle development. These findings reveal a novel molecular mechanism by which the OsNLP3/4-OsRFL-OsMOC1 regulatory module integrates N availability to promote panicle architecture in rice.

## Results

### OsNLP3/4 positively regulate the rice panicle architecture

Our previous study identified OsNLP3/4 as key players in rice yield and NUE (Wu *et al*., 2021; Zhang *et al*., 2022). To confirm whether panicle architecture is a direct contributor to the increased yield, we observed panicle morphology in the wild type (WT), *OsNLP3*/*4* knockout mutants (*nlp3-2*, *nlp3-3*, *nlp4-1*, *nlp4-2*) and their overexpression plants (OE3-1, OE3-10, OE4-1, OE4-9) and found that the *OsNLP3*/*4*- OE plants exhibited larger-panicle phenotypes than other genotypes (Fig. 1a, f).

**Fig. 1.**
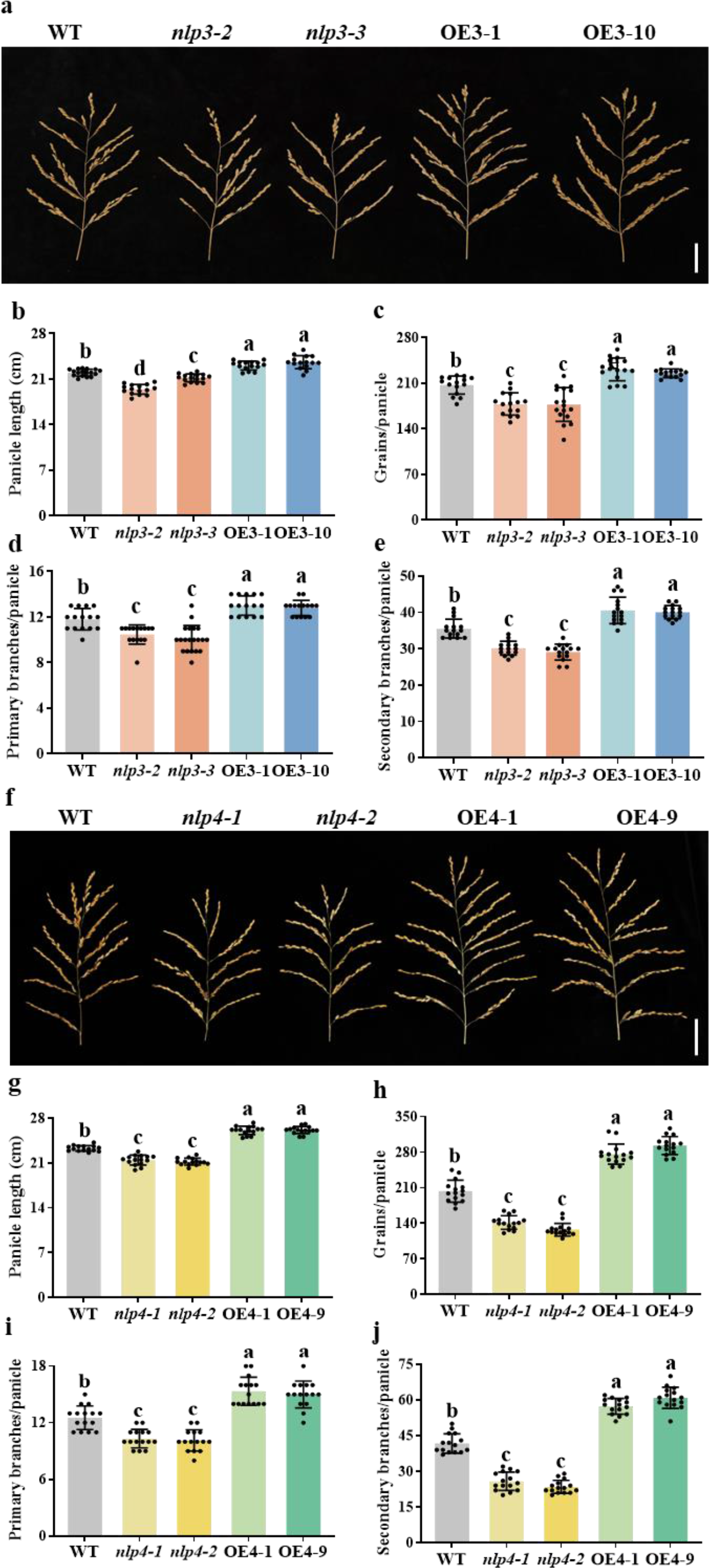
Panicle architecture of different *OsNLP3*/*4* genotypes under normal conditions. (**a**) Mature panicle morphology of the WT, *nlp3* mutants (*nlp3-2* and *nlp3-3*), and *OsNLP3* overexpression lines (OE3-1 and OE3-10) under normal conditions in Lingshui (December 2020 to April 2021). Bar = 5 cm. (**b-e**) Panicle architecture-related traits of the WT and different *OsNLP3* genotypes in (a) under normal conditions, including panicle length (b), number of grains per panicle (c), number of primary branches per panicle (d), and number of secondary branches per panicle (e). Values are means ± SD (n = 15). Different letters indicate significant differences by one-way ANOVA with Tukey’s test (*P* < 0.05). (**f**) Mature panicle morphology of the WT, *nlp4* mutants (*nlp4-1* and *nlp4-2*), and *OsNLP4* overexpression lines (OE4-1 and OE4-9) under normal conditions in Bengbu (May 2021 to October 2021). Bar = 5 cm. (**g-j**) Panicle architecture-related traits of the WT and different *OsNLP4* genotypes in (f) under normal conditions, including panicle length (g), number of grains per panicle (h), number of primary branches per panicle (i), and number of secondary branches per panicle (j). Values are means ± SD (n = 15). Different letters indicate significant differences by one-way ANOVA with Tukey’s test (*P* < 0.05).

Compared with the WT, the *OsNLP3*-OE plants showed an average 6.10% increase in panicle length, accompanied by a significant increase of 10.33% in grain number per panicle, whereas the *nlp3* mutants displayed a 7.65% decrease in panicle length and 14.87% of grain number per panicle (Fig. 1b, c). The number of primary and secondary panicle branches, the main factors responsible for the differences in grain number per panicle, increased by 10.38% and 14.01% in the *OsNLP3*-OE plants and decreased by 11.97% and 16.70% in the *nlp3* mutants, respectively, compared with those of the WT (Fig. 1d, e).

*OsNLP4* played a similar but stronger role in panicle architecture than *OsNLP3*. The *OsNLP4*-OE plants exhibited a significant increase in panicle length and grain number per panicle by 12.16% and 40.34%, respectively, whereas the *nlp4* mutants displayed a sharp decrease by 8.43% and 33.76% compared with the WT (Fig. 1g, h). Consistently, the number of both primary and secondary branches increased significantly in the *OsNLP4*-OE plants by 20.82% and 41.58%, respectively, while the *nlp4* mutants decreased by 18.32% and 40.94% compared with those in the WT (Fig. 1i, j). Together, these results clearly show that OsNLP3/4 are positive regulators of rice panicle architecture.

### OsNLP3/4-promoted panicle architecture positively correlates with N availability

The ability of N to influence panicle development and induce the nuclear localization of OsNLP3/4 prompted us to test whether OsNLP3/4-regulated panicle development responds to N availability. We explored the panicle architecture of different *OsNLP3*/*4* genotypes under different N concentrations (Fig. 2a, f). Under low nitrogen (LN) conditions, the *OsNLP3*-OE plants exhibited an average increase of 12.43% in panicle length and 13.12% in grain number per panicle compared with the WT, whereas the *nlp3* mutants showed a decrease of 10.35% in panicle length and 18.28% in grain number per panicle. Under normal N (NN) conditions, the *OsNLP3*- OE plants showed an average increase of 13.78% in panicle length and 18.79% in grain number per panicle compared with the WT, whereas the *nlp3* mutants exhibited a decrease of 11.42% in panicle length and 21.27% in grain number per panicle.

**Fig. 2.**
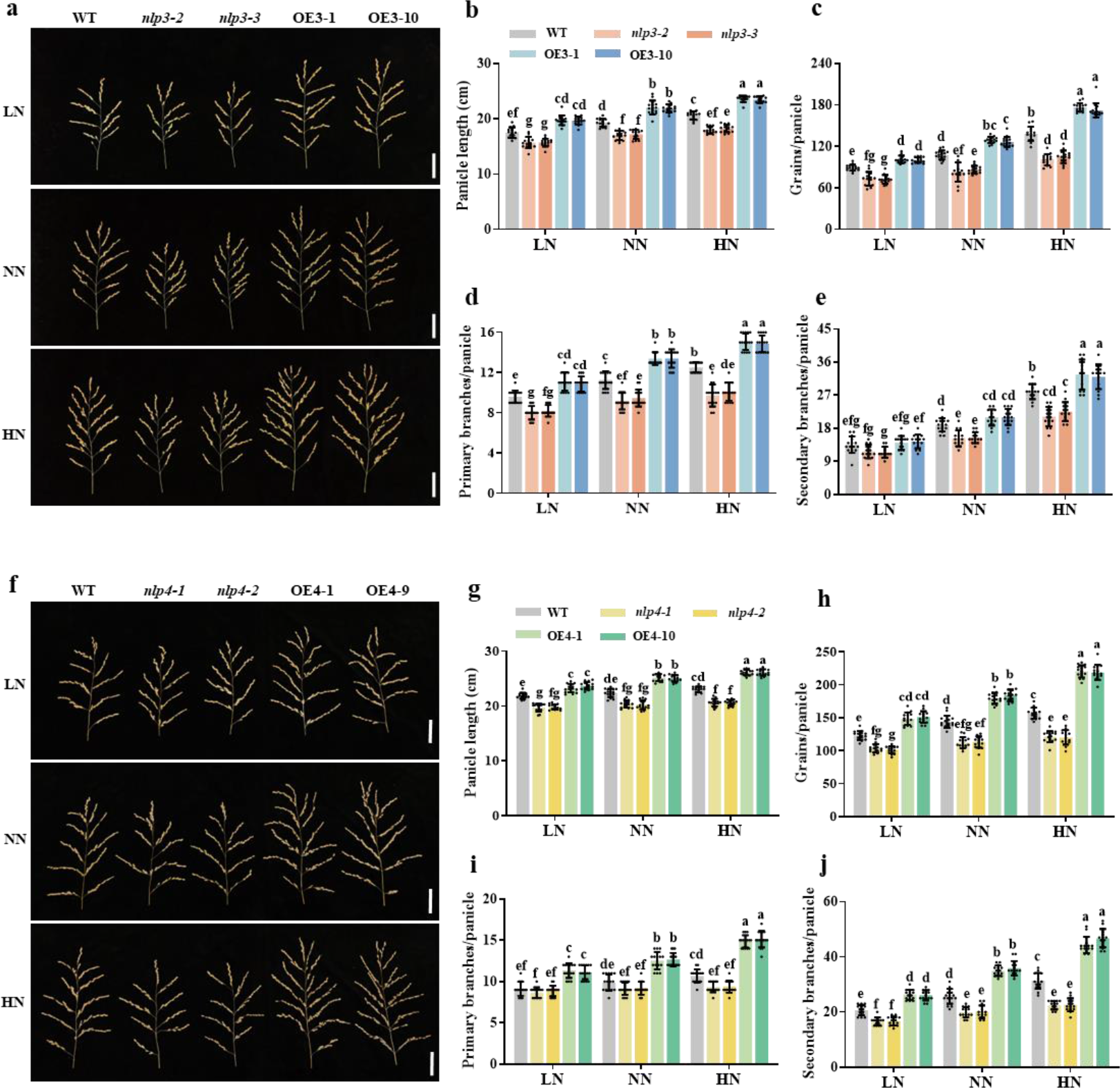
Panicle architecture of different *OsNLP3*/4 genotypes under different nitrogen conditions. (**a**) Mature panicle morphology of the WT, *nlp3* mutants (*nlp3-2* and *nlp3-3*), and *OsNLP3* overexpression lines (OE3-1 and OE3-10) under low nitrogen (LN), normal nitrogen (NN) and high nitrogen (HN) conditions in Hefei (May 2022 to October 2022). Bars = 5 cm. (**b-e**) Panicle architecture-related traits of the WT and different *OsNLP3* genotypes in (a) under different nitrogen conditions, including panicle length (b), number of grains per panicle (c), number of primary branches per panicle (d), and number of secondary branches per panicle (e). Values are means ± SD (n = 15). Different letters indicate significant differences by one-way ANOVA with Tukey’s test (*P* < 0.05). (**f**) Mature panicle morphology of the WT, *nlp4* mutants (*nlp4-1* and *nlp4-2*), and *OsNLP4* overexpression lines (OE4-1 and OE4-9) under low nitrogen (LN), normal nitrogen (NN) and high nitrogen (HN) conditions in Changxing (May 2021 to October 2021). Bars = 5 cm. (**g-j**) Panicle architecture-related traits of the WT and different *OsNLP4* genotypes in (f) under different nitrogen conditions, including panicle length (g), number of grains per panicle (h), number of primary branches per panicle (i), and number of secondary branches per panicle (j). Values are means ± SD (n = 15). Different letters indicate significant differences by one-way ANOVA with Tukey’s test (*P* < 0.05).

Under high N (HN) conditions, the *OsNLP3*-OE plants displayed a significant increase of 14.25% in panicle length and 26.18% in grain number per panicle compared with the WT, whereas the *nlp3* mutants showed a sharp decrease of 12.02% in panicle length and 25.53% in grain number per panicle (Fig. 2b, c, Table S1). The number of primary and secondary branches responded to N concentrations in a similar trend as grain number per panicle in the three genotypes (Fig. 2d, e, Table S1).

Similarly, the panicle length, grain number per panicle, primary and secondary branch numbers of the *OsNLP4*-OE plants increased significantly from 7.38%, 21.79%, 22.99% and 27.02% under LN conditions to 13.27%, 40.00%, 40.62% and 45.63% under HN conditions compared to the WT, respectively, while those parameters of *nlp4* mutants decreased dramatically from 9.57%, 16.68%, 4.01% and 18.93% under LN conditions to 10.53%, 22.99%, 12.19% and 28.04% under HN conditions (Fig. 2g-j, Table S1). Similar panicle architecture results were reproduced in multiple independent experimental replicates, as shown in Fig. S1. Together, these results demonstrate that the OsNLP3/4-promoted panicle architecture is positively correlated with the levels of N supply.

### OsNLP3/4 affect the development of inflorescence meristem

The size and number of inflorescence meristems (IMs) are directly correlated with rice panicle architecture. To contrast the panicle development of different *OsNLP3*/*4* genotypes, we observed the morphology of the young inflorescences during the transition from the vegetative to reproductive stage and primary branching stage by scanning electron microscopy (SEM) (Fig. 3). The IM sizes of the *OsNLP3*- OE and *OsNLP4*-OE plants were significantly larger than those of the WT (Fig. 3d-f), whereas the *nlp3* and *nlp4* mutants exhibited obviously smaller IM sizes (Fig. 3b-c, f). Further observations showed that the *OsNLP3*-OE and *OsNLP4*-OE plants had more primary branching meristems (PBMs), which increased by 22.76% and 23.73% compared with the WT plants, respectively (Fig. 3i-j, m-n). Conversely, the *nlp3* and *nlp4* mutants showed 33.28% and 25.24% fewer PBMs than the WT inflorescences (Fig. 3h, j, l, n). Together, these results indicate that *OsNLP3*/*4* are positive regulators of IM size, consistent with our panicle data (Fig. 1 and 2).

**Fig. 3.**
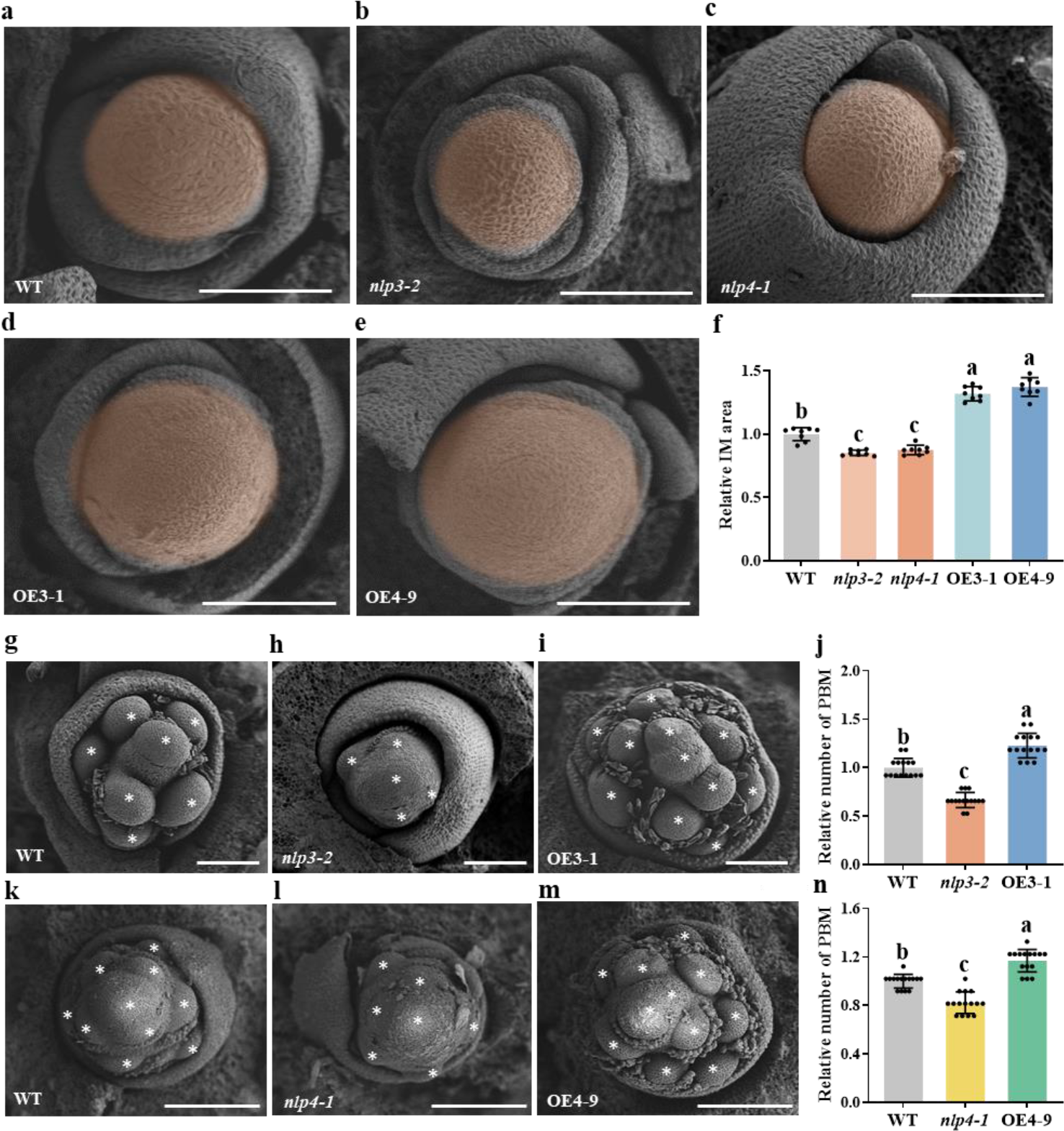
Scanning electron microscope (SEM) analysis of inflorescence meristem (IM) size and primary branch meristem (PBM) number in *OsNLP3*/*4* genotypes. (a-e) SEM images showing IM size of the WT (a), *nlp3-2* (b), *nlp4-1* (c), OE3-1 (d) and OE4-9 (e). The orange areas indicate IM. Bars = 100 μm. **(f)** Average IM area of the WT, *nlp3-2*, *nlp4-1*, OE3-1 and OE4-9. Values are means ± SD relative to the WT, which is set to 1 (n = 8). Different letters indicate significant differences by one-way ANOVA with Tukey’s test (*P* < 0.05). **(g-i, k-m)** SEM images showing PBM number in the WT (g), *nlp3-2* (h), OE3-1 (i), and the WT (k), *nlp4*-*1* (l), OE4-9 (m). The asterisks indicate PBMs. Bars = 100 μm. (**j, n**) Average PBM numbers of the WT, *nlp3-2*, OE3-1 (j) and the WT, *nlp4*-*1*, OE4-9 (n). Values are means ± SD relative to the WT, which is set to 1 (n = 15). Different letters indicate significant differences by one-way ANOVA with Tukey’s test (*P* < 0.05).

### OsNLP3/4 upregulates *OsRFL* expression to increase panicle size

To explore the mechanism by which OsNLP3/4 promote panicle architecture, we performed transcriptomic analyses of developing inflorescence and identified *OsRFL* as a candidate target of OsNLP3/4, which is a positive regulator of panicle architecture by promoting IM development in rice (Huang *et al*., 2021; Ikeda-Kawakatsu *et al*., 2012). *OsRFL* expression was promoted in the *OsNLP3*-OE and *OsNLP4*-OE plants and suppressed in the *nlp3* and *nlp4* mutants (Fig. 4a-b), and its transcript levels were positively correlated with N concentrations (Fig. S2), suggesting that *OsRFL* may be a target gene of OsNLP3/4.

**Fig. 4.**
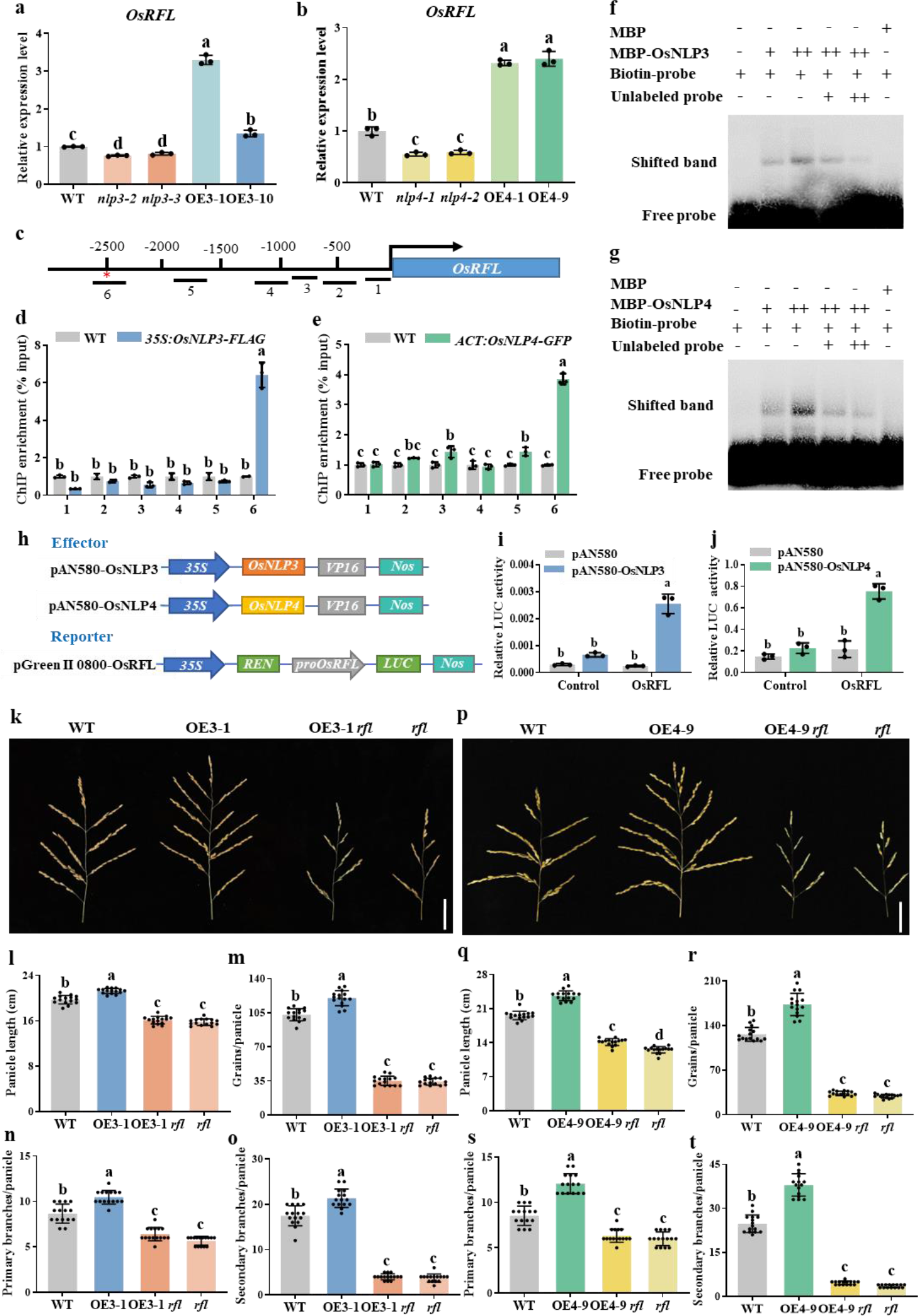
Both OsNLP3 and OsNLP4 activate *OsRFL* by binding to its promoter. (a-b) Relative expression of *OsRFL* in different *OsNLP3* (a) and *OsNLP4* genotypes (b), normalized to *OsACTIN1* at the branch meristem formation stage (0.5 cm young panicle). Values are means ± SD relative to WT, which is set to 1 (n = 3). Different letters indicate significant differences by one-way ANOVA with Tukey’s test (*P* < 0.05). (**c-e**) ChIP assays. (c) The diagram depicts the regions used for ChIP-qPCR analysis. The red asterisk indicates the common binding region of OsNLP3 and OsNLP4. The enrichments of the *OsRFL* promoter analyzed by ChIP-qPCR using extracts from young panicles (0.5-3 cm) of the *35S:OsNLP3-FLAG* transgenic rice plants (d) and *OsACTIN1pro:OsNLP4*-*GFP* transgenic rice (e) were normalized to input DNA. Values are means ± SD relative to the WT, which is set to 1 (n = 3). Different letters indicate significant differences by one-way ANOVA with Tukey’s test (*P* < 0.05). (**f-g**) EMSA to detect OsNLP3 (f) and OsNLP4 (g) binding to the *OsRFL* promoter region containing the NRE-like *cis*-element. The shifted band and free probes are indicated. (**h-j**) Dual-luciferase assays. (h) The diagram depicts the effectors and reporter used in the dual-luciferase assays. The effector genes *OsNLP3* and *OsNLP4* were under the control of the *CaMV 35S* promoter. One thousand bp fragment containing NRE-like *cis*-elements from the promoter of *OsRFL* were fused to the *LUC* gene as a reporter. (i-j) Firefly LUC and REN luciferase activities were detected by transient dual-luciferase reporter assays, and the LUC:REN ratio was calculated. REN activity was used as an internal control. Values are means ± SD (n = 3). Different letters indicate significant differences by one-way ANOVA with Tukey’s test (*P* < 0.05). (**k**) Mature panicle morphology of the WT, *OsNLP3* overexpression lines (OE3-1), *OsNLP3* overexpression lines in the *rfl* mutant background (OE3-1 *rfl*), and the *rfl* mutant grown under normal conditions in Bengbu (May 2022 to October 2022). Bar = 5 cm. (**l-o**) Panicle architecture-related traits of four genotypes in (k) under normal conditions, including panicle length (l), number of grains per panicle (m), number of primary branches per panicle (n), and number of secondary branches per panicle (o). Values are means ± SD (n = 15). Different letters indicate significant differences by one-way ANOVA with Tukey’s test (*P* < 0.05). (**p**) Mature panicle morphology of the WT, *OsNLP4* overexpression lines (OE4-9), *OsNLP4* overexpression lines under the *rfl* mutant background (OE4-9 *rfl*), and the *rfl* mutant grown under normal conditions in Lingshui (December 2021 to April 2022). Bar = 5 cm. (**q-t**) Panicle architecture-related traits of four genotypes in (p) under normal conditions, including panicle length (q), number of grains per panicle (r), number of primary branches per panicle (s), and number of secondary branches per panicle (t). Values are means ± SD (n = 15). Different letters indicate significant differences by one-way ANOVA with Tukey’s test (*P* < 0.05).

We performed a *cis* element search in the promoter of *OsRFL* and found putative nitrate-responsive *cis*-elements (NREs), the potential binding sites of OsNLP3/4.

Chromatin immunoprecipitation (ChIP) assays showed that the *cis*6 portion of the *OsRFL* promoter harboring NRE was significantly enriched, confirming that OsNLP3/4 bind to the *OsRFL* promoter *in vivo* (Fig. 4c-e). These binding events were further confirmed by electrophoretic mobility shift assays (EMSAs) (Fig. 4f, g).

Furthermore, we conducted a dual-luciferase (LUC) reporter assay to further verify the transcriptional activation of *OsRFL* by OsNLP3/4 in rice protoplasts. Strong LUC activities were shown when the effector construct *35S-OsNLP3* or *35S-OsNLP4* was cotransfected with the reporter construct *OsRFLpro:LUC* (Fig. 4h-j), indicating that OsNLP3/4 transcriptionally activate the expression of *OsRFL*.

To genetically confirm the relationship between *OsNLP3*/*4* and *OsRFL*, we performed genetic analyses of the *rfl*, OE3-1 *rfl* and OE4-9 *rfl* mutants generated by editing *OsRFL* in the WT, *OsNLP3*-OE and *OsNLP4*-OE backgrounds using CRISPR- Cas9 techniques (Fig. S3). Panicle development was severely compromised in the *rfl* mutants, with an average reduction of 50% in panicle length, 70% in primary branches and 80% in secondary branches compared with the WT, resulting in a sharp 85% reduction in grain number per panicle (Fig. 4k-t), consistent with previous reports (Huang *et al*., 2021; Ikeda-Kawakatsu *et al*., 2012). The panicle architecture of the OE3-1 *rfl* and OE4-9 *rfl* mutants was similar to that of *rfl* (Fig. 4k-t), indicating that *OsNLP3*/*4* are in a common genetic pathway and epistatic to *OsRFL*. Taken together, these results clearly demonstrate that OsNLP3/4 positively regulate *OsRFL* to control panicle architecture.

### *OsMOC1* is a direct target gene of OsRFL

Although OsRFL is known as a key transcription factor regulating panicle development, its downstream regulatory network is largely unknown until recently several target genes of OsRFL have been identified (Miao et al., 2022), including *OsSPL7*, *OsSPL14*, *OsSPL17*, *NECK LEAF1* (*NL1*), and *OSH6*. To identify more downstream target genes regulated by the OsNLP3/4-OsRFL module, we examined the transcript levels of several marker genes involved in IM maintenance and specification, including *OsLAX1*, *OsLAX2*, *OsMOC1*, and *OsMOC3*, in different *OsNLP3*/*4* genotypes. The expression of these genes was upregulated in *OsNLP3*-OE and *OsNLP4*-OE plants but downregulated in *nlp3* and *nlp4* mutants (Fig. S4), suggesting that the OsNLP3/4-OsRFL module may affect panicle architecture by regulating these genes.

*OsMOC1* showed similar expression patterns as *OsRFL* among all genotypes of *OsNLP3*/*4*, being upregulated in the *OsNLP3*-OE and *OsNLP4*-OE plants and downregulated in the *nlp3* and *nlp4* mutants under different N conditions (Fig. 5a, b; Fig. S2). Meanwhile, *OsMOC1* expression was severely inhibited in the *rfl* mutants (Fig. 5c). In addition, the *OsMOC1* promoter contains the WNNN (CCANTG) G/TNNNW *cis*-element recognized by LEAFY (LFY), a homolog of OsRFL in *Arabidopsis* (Jin et al., 2021; Rao *et al*., 2008), suggesting that OsRFL may directly bind to the *OsMOC1* promoter to regulate its expression. EMSA confirmed the binding of OsRFL to the *OsMOC1* promoter region containing the WNNN (CCANTG) G/TNNNW *cis*-element *in vitro* (Fig. 5d). Subsequent transcriptional activation assays showed that OsRFL activated the expression of the *LUC* gene linked to the *OsMOC1* promoter (Fig. 5e, f). These results demonstrate that OsRFL binds to the promoter of *OsMOC1* and activates its expression.

**Fig. 5.**
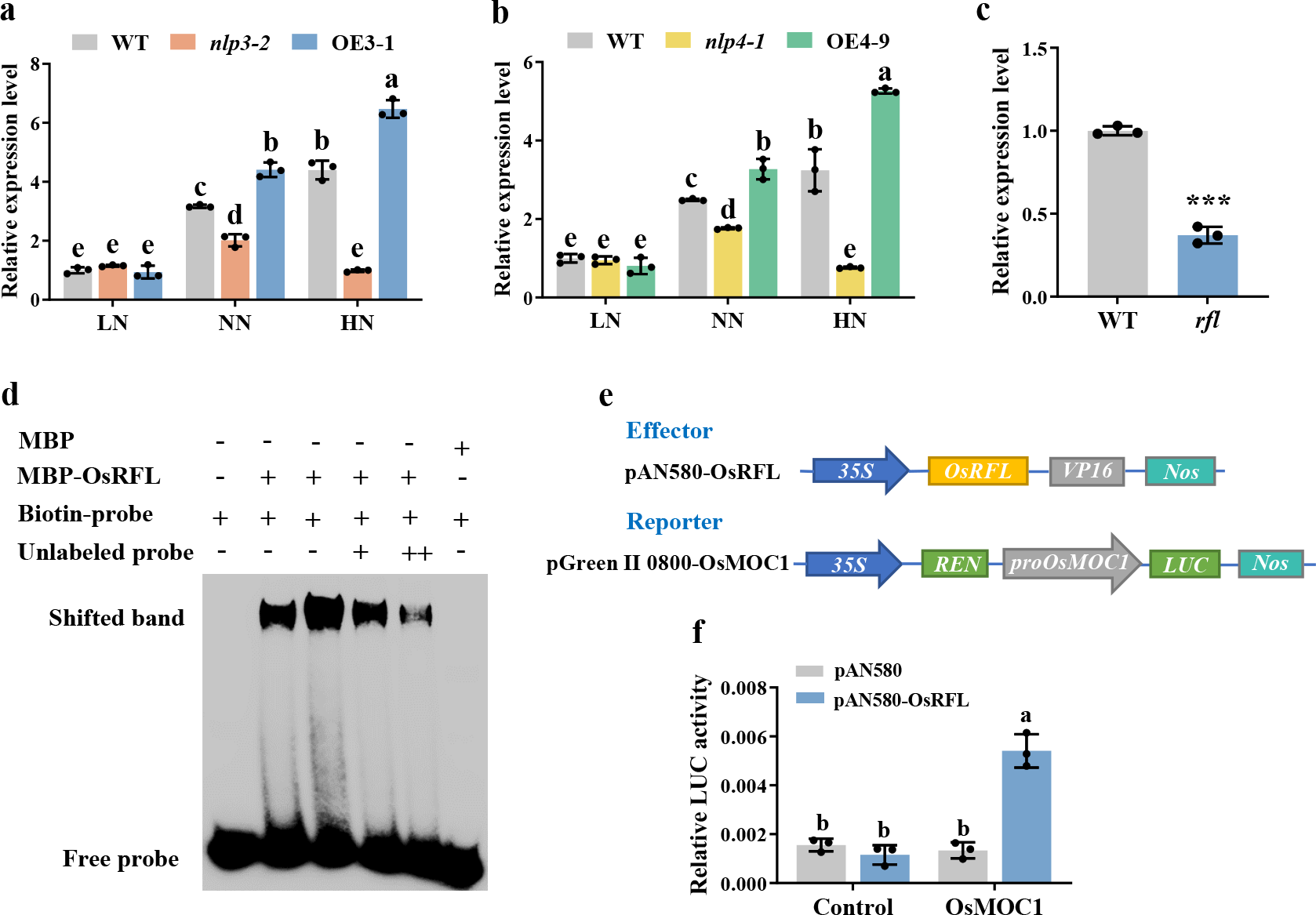
OsRFL activates *OsMOC1* by binding to its promoter. (**a-b**) Relative expression of *OsMOC1* in different *OsNLP3* (a) and *OsNLP4* genotypes (b) grown in vermiculite pots under 0.5 mM nitrate (LN), 1 mM nitrate (NN), and 5 mM nitrate (HN) conditions. The expression levels are normalized to *OsACTIN1* at the branch meristem formation stage (0.5 cm young panicle). Values are means ± SD relative to WT under the LN conditions, which is set to 1 (n = 3). Different letters indicate significant differences by one-way ANOVA with Tukey’s test (*P* < 0.05). (c) Relative expression of *OsMOC1* in *rfl*, normalized to the rice *OsACTIN1* gene at the branch meristem formation stage (0.5 cm young panicle). Values are means ± SD relative to the WT, which is set to 1 (n = 3). Asterisks indicate significant differences by Student’s *t* test (\**P* < 0.05, *\*\*P* < 0.01, *\*\*\*P* < 0.001). (d) EMSA to detect OsRFL binding to the *OsMOC1* promoter region containing the WNNN (CCANTG) G/TNNNW *cis*-element. The shifted band and free probe are indicated. (**e-f**) Dual-luciferase assays. (e) The diagram depicts the effector and reporter used in the dual-luciferase assays. The effector gene *OsRFL* was under the control of the *CaMV 35S* promoter. One thousand bp fragment containing the WNNN (CCANTG) G/TNNNW *cis*-element from the promoter of *OsMOC1* was fused to the *LUC* gene as a reporter. (f) The firefly LUC and REN luciferase activities were detected by transient dual-luciferase reporter assays, and the LUC:REN ratio was calculated. REN activity was used as an internal control. Values are means ± SD (n = 3). Different letters indicate significant differences by one-way ANOVA with Tukey’s test (*P* < 0.05).

## Discussion

Panicle architecture is an important agronomic trait and a major contributor to rice yield (Xing and Zhang, 2010). Although a number of genes involved in panicle development have been identified in recent years (Zhang and Yuan, 2014), the molecular mechanisms underlying the response of panicle architecture to nutrients remain unclear. In this study, we revealed a novel function of OsNLP3/4 in the regulation of panicle architecture in rice. Using knockout mutants and overexpression lines, we demonstrated with strong genetic evidence that OsNLP3/4 are positive regulators of rice panicle architecture (Fig. 1) and that the OsNLP3/4-regulated panicle architecture positively correlates with N concentrations (Figs. 2, S1).

Importantly, we revealed that the N-responsive OsNLP3/4 directly upregulate the expression of *OsRFL* (Fig. 4), encoding an important regulator of IM development (Huang *et al*., 2021; Ikeda-Kawakatsu *et al*., 2012), and OsRFL upregulates *OsMOC1* (Fig. 5), coding for a GRAS family transcription factor that regulates IM formation and development, which was strongly supported by our IM anatomic data (Fig. 3).

Therefore, this study revealed a novel molecular mechanism by which the N-responsive OsNLP3/4-OsRFL module integrates N availability and positively regulates the expression of *OsMOC1* to promote rice IM development, thus improving panicle architecture. A working model is proposed in Fig. 6.

**Fig. 6.**
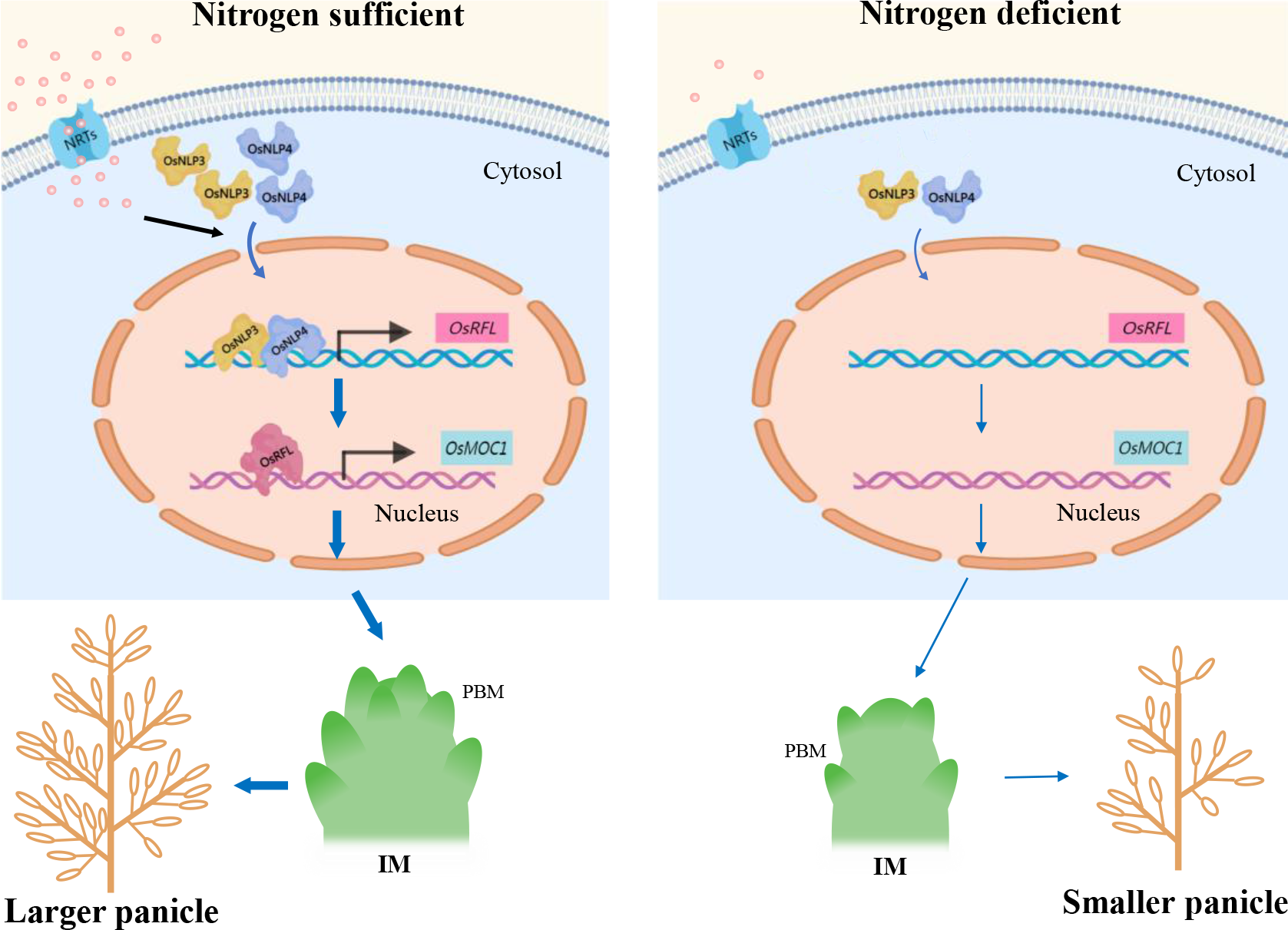
A working model proposed for the OsNLP3/4-OsRFL-OsMOC1 module that integrates N availability to regulate panicle architecture. Under N-sufficient conditions (left panel), most of the OsNLP3/4 proteins are translocated from the cytoplasm to the nucleus and strongly promote *OsRFL* expression, which subsequently upregulates *OsMOC1* to control IM development and ultimately improves panicle architecture. Under N-deficient conditions (right panel), the majority of OsNLP3/4 proteins are localized in the cytoplasm but rarely in the nucleus and cannot fully activate the expression of the *OsRFL*-*OsMOC1* module, ultimately producing small panicles. The thicknesses of the arrows schematically represent the relative strength of regulation. The small pink circles around NRTs outside or inside cytoplasm represent nitrate.

### A novel function of OsNLP3/4 in panicle development

NLPs are pivotal transcription factors for early N signaling, N fixation and cold stress responses and act as receptors of nitrate, as recently reported for AtNLP7 (Ding et al., 2022; Jiang *et al*., 2021; Liu *et al*., 2022; Mu and Luo, 2019). However, the regulation of NLPs in development has rarely been studied. AtNLP8 promotes seed germination in a nitrate-dependent manner by directly binding to the promoter of *CYTOCHROME P707A2* (*CYP707A2*) to reduce ABA levels (Yan *et al*., 2016). In this study, we discovered that OsNLP3/4, which were previously reported to promote rice yield and NUE (Wu *et al*., 2021; Zhang *et al*., 2022), improved the panicle architecture by upregulating *OsRFL* expression and thereby IM development, increasing the size of IMs and the number of PBMs, resulting in a significant increase in panicle length, primary and secondary branches, and grains per panicle (Figs. 1, 3).

*OsRFL* is a target gene of OsNLP3/4 (Fig. 4a, b). OsNLP3/4 directly bound to the NRE elements in the *OsRFL* promoter to transcriptionally activate its expression (Fig. 4c-j). The OE3-1 *rfl* and OE4-9 *rfl* mutants exhibited similar aberrant panicle architecture as *rfl* (Fig. 4k-t), providing strong genetic evidence that *OsNLP3*/*4* are epistatic to *OsRFL*.

LFY, the homolog of OsRFL in *Arabidopsis*, is a pioneer transcription factor capable of regulating multiple genes, such as the expression of *APETALA1* (*AP1*), to activate IM fate (Jin *et al*., 2021). However, the downstream target genes of OsRFL regulating IM development remain largely unclear. Recently, by combining RNA-seq and ChIP-seq assays, Miao et al. found that OsRFL can directly regulate *OsSPL7*, *OsSPL14*, *OsSPL17*, *NL1*, and *OSH6* to control panicle architecture (Miao *et al*., 2022), while omics data suggest that there are still more genes regulated by OsRFL yet to be identified. *OsMOC1*, encoding a GRAS family transcription factor, also regulates IM formation and development. The *moc1* mutants exhibit abnormal panicles similar to *rfl* (Li et al., 2003; Zhang *et al*., 2020). Our molecular and biochemical experiments showed that OsRFL could activate *OsMOC1* expression by binding to the WNNN (CCANTG) G/TNNNW *cis*-element within its promoter (Fig. 5c-f), suggesting that *OsMOC1* is a target gene of OsRFL that regulates panicle development. Moreover, *OsMOC1* and *OsRFL* showed similar expression patterns in the different genotypes of *OsNLP3*/*4*, suggesting that the OsRFL-OsMOC1 module is under the control of *OsNLP3*/*4* (Figs. 4a, 4b, 5a, 5b, S2, S4). Therefore, we reveal the novel OsNLP3/4-*OsRFL*-*OsMOC1* module that promotes panicle development.

### The OsNLP3/4-OsRFL module integrates N availability to promote panicle architecture

As one of the essential macronutrients for plants, increased availability of N promotes rice panicle development and panicle grain number, but the underlying regulatory mechanism remains unclear (Ding *et al*., 2014; Luo *et al*., 2020). We found that the *OsNLP3*/*4*-regulated panicle architecture was positively correlated with the level of N supply. Grain number per panicle increased from 13.12% and 21.79% in *OsNLP3*/*4* OE plants under LN conditions to 26.18% and 40.00% under HN conditions compared with those in WT. Likewise, panicle length and primary and secondary branches also showed similar trends (Fig. 2, Table S1).

NLP proteins are known for their N-responsive nucleocytosolic shuttling, leading to changes in downstream events. For example, the regulation of AtNLP7 for root and shoot growth is partly mediated by the nitrate-induced nuclear accumulation that triggers transcriptional reprogramming (Liu *et al*., 2017). MtNLP1 shuttles from the cytosol to the nucleus in response to nitrate, thereby interacting with MtNIN to inhibit nodule formation under N-sufficient conditions (Lin *et al*., 2018). In crops, ZmNLP6, ZmNLP8, and HvNLP2 regulate NUE through a nitrate-mediated nuclear retention mechanism (Cao *et al*., 2017; Gao *et al*., 2021). It is also well known that the nuclear accumulation of OsNLP3/4 positively correlates with the availability of N (Wu *et al*., 2021; Zhang *et al*., 2022). In this study, we further showed that the expression of *OsRFL* and *OsMOC1* was also positively correlated with N supply (Figs. 5a-b, S2).

Together, these findings strongly support that N regulates rice panicle architecture through the OsNLP3/4-OsRFL-OsMOC1 module (Fig. 6).

### OsNLP3/4 performs overlapping but differential roles in panicle architecture regulation

Both the *OsNLP3*-OE and *OsNLP4*-OE plants exhibited larger-panicle phenotypes, and the *nlp3* and *nlp4* mutants showed similar abnormal panicle architectures with shortened rachis, reduced branching and grain, due to regulation of the common OsRFL-OsMOC1 module (Figs. 1-5, S1, Table S1), suggesting overlapping functions of OsNLP3/4 in rice panicle architecture regulation. However, *OsNLP4* played a stronger role in panicle architecture than *OsNLP3* (Figs. 1, 2, S1, Table S1). For example, compared with the WT, the panicle length, grain number per panicle, primary and secondary branch numbers increased by 6.10%, 10.33%, 10.38% and 14.01% in the *OsNLP3-OE* plants, and by 12.16%, 40.34%, 20.82%, and 41.58% in the *OsNLP4-OE* plants, respectively (Fig. 1). One possible reason is that the nuclear localization of OsNLP4 was induced by both nitrate and ammonium, while that of OsNLP3 was only induced by nitrate (Wu *et al*., 2021; Zhang *et al*., 2022). In addition, *OsNLP4* was more biased toward the regulation of secondary branching than *OsNLP3*, as reflected by the larger gap in secondary branch numbers among the *OsNLP4* genotypes (Figs. 1e, 1j, 2e, 2j, S1e, S1j, Table S1), which further indicates the functional differentiation of OsNLP3 and OsNLP4 in the regulation of panicle architecture.

Multiple interactions between different NLP transcription factors are mediated through their PB1 domains (Guan et al., 2017; Konishi and Yanagisawa, 2019).

Homodimerizations of NLPs in *Arabidopsis* are not required for the transcriptional activation of nitrate-responsive genes, but are beneficial to promote the high expression of these genes (Konishi and Yanagisawa, 2019). Additional studies are needed to dissect whether OsNLP3 and OsNLP4 interact to cooperatively regulate panicle architecture. Moreover, even under LN conditions, the panicle architecture of the *nlp3* and *nlp4* mutants was not as severe as that of *rfl* (Figs. 2, 4), suggesting that there may be other proteins whose localization is not affected by N, such as OsNLP1 (Alfatih et al., 2020), undertaking a similar function to OsNLP3/4. Together, these results demonstrate that OsNLP3/4 have both conserved and differentiated functions in rice panicle architecture regulation.

In conclusion, our study identified the novel OsNLP3/4-OsRFL-OsMOC1 module that integrates N availability to positively promote the development of rice panicles, revealing a molecular mechanism for N-promoted rice panicle architecture and opening a new window for the improvement of grain yield via panicle architecture.

## Methods

### Plant materials

Wild-type rice (*Oryza sativa* L.) *japonica* variety Zhonghua11 (ZH11) was used in this study. The loss-of-function mutants (*nlp3-2*, *nlp3-3*, *nlp4-1*, *nlp4-2*) and overexpression lines of *OsNLP3* (OE3-1, OE3-10) and *OsNLP4* (OE4-1, OE4-9) were reported previously (Wu *et al*., 2021; Zhang *et al*., 2022). Mutants *rfl*, OE3-1 *rfl* and OE4-9 *rfl* were generated in the WT, *OsNLP3* overexpression and *OsNLP4* overexpression backgrounds, respectively, generated by Wimi Biotechnology Co., Ltd. (Changzhou, China) (http://www.wimibio.com/) via the CRISPR/Cas9 technique (Liu et al., 2020). For ChIP assays, the *35S:OsNLP3-FLAG* and *OsACTIN1pro:OsNLP4- GFP* transgenic plants were reported previously (Hu et al., 2019; Wu *et al*., 2021).

### Plant growth conditions

### Plants in the greenhouse

Seeds were washed with distilled water and incubated at 37 ℃ for 3 days. Germinated seeds were transferred to soil for regular water and fertilizer management. The provided growth conditions were kept at a 16-h light (30 °C)/8-h dark (28 °C) cycle.

### Plants in the field

Field tests were carried out in paddy fields under natural growth conditions and different N conditions during 2019-2022 at five experimental locations: Bengbu (Anhui Province, China), Lingshui (Hainan Province, China), Hefei (Anhui Province, China), Chengdu (Sichuan Province, China), and Changxing (Zhejiang Province, China).

Bengbu field trials were performed in 2021 (May to October) for the WT, *nlp4* mutants (*nlp4-1* and *nlp4-2*), and *OsNLP4* overexpression lines (OE4-1 and OE4-9) and in 2022 (May to October) for the WT, OE3-1, *rfl*, and OE3-1 *rfl* plants under normal conditions. Urea was used as the N source at 185 kg N/ha. The plants were transplanted in 8 rows × 10 plants for each plot (3.6 m^2^), and four replicates were used for each treatment.

The Lingshui field trials were performed from December 2020 to April 2021 for the WT, *nlp3* mutants (*nlp3-2* and *nlp3-3*), and *OsNLP3* overexpression lines (OE3-1 and OE3-10) and from December 2021 to April 2022 for the WT, OE4-9, *rfl*, and OE4-9 *rfl* plants under normal conditions. Urea was used as the N source at 150 kg N/ha. The plants were transplanted in 10 rows×10 plants for each plot (4 m^2^), and four replicates were used for each treatment.

The Hefei field trial was performed in 2022 (May to October) for the WT, *nlp3* mutants (*nlp3-2* and *nlp3-3*), and *OsNLP3* overexpression lines (OE3-1 and OE3-10) under different N conditions. Urea was used as the N source, with 80 kg N/ha for low N, 160 kg N/ha for normal N and 320 kg N/ha for high N. The plants were transplanted in 8 rows×10 plants for each plot (3.4 m^2^), and four replicates were used for each treatment.

The Chengdu field trial was performed in 2019 (April to September) for the WT, *nlp3* mutants (*nlp3-2* and *nlp3-3*), and *OsNLP3* overexpression lines (OE3-1 and OE3-10) under different N conditions. Urea was used as the N source, with 80 kg N/ha for low N, 200 kg N/ha for normal N and 500 kg N/ha for high N. The plants were transplanted in 10 rows×20 plants for each plot (8 m^2^), and four replicates were used for each treatment.

The Changxing field trials were performed in 2019 (May to October) and 2021 (May to October) for the WT, *nlp4* mutants (*nlp4-1* and *nlp4-2*), and *OsNLP4* overexpression lines (OE4-1 and OE4-9) under different N conditions. Urea was used as the N source, with 90 kg N/ha for low N, 180 kg N/ha for normal N and 360 kg N/ha for high N. The plants were transplanted in 8 rows×10 plants for each plot (3.4 m^2^), and four replicates were used for each treatment.

### Measurement of panicle architecture

Panicles of mature rice were harvested, and representative panicles were imaged by a digital camera (Nikon, D700). Main panicles from the mature plants were used to analyze panicle architecture.

### Scanning electron microscopy (SEM) analysis

Approximately two-month-old seedlings grown in soil in a greenhouse were taken for scanning electron microscopy analysis. Scanning electron microscopy of inflorescence meristem size and primary branch primordia was performed as described previously with modifications (Ikeda *et al*., 2005). After fixation in 2.5% glutaraldehyde (dissolved in 0.1 mol/L PBS, pH 7.2) at 4 ℃ overnight and dehydration in a graded ethanol series, the samples were replaced with acetone and dried with a critical-point dryer (Emitech, K850) immediately. Subsequently, we sputtered the samples with gold (Leica, EM ACE200) and observed them with a scanning electron microscope (Zeiss, Genimi SEM 500). Inflorescence meristem sizes were calculated by ImageJ software.

### RNA extraction and qRT-PCR

Total RNA was extracted using TRIzol reagent (TransGen, ET101-01) from young panicles (0.5 cm, in the branch primordium stage) of plants grown in soil in a greenhouse. Two micrograms of total RNA were used to synthesize cDNA using the Easyscript One-step gDNA Removal and cDNA Synthesis SuperMix reagent kit (TransGen, AE311-01). We used the SYBR Premix Ex Taq II reagent kit (Vazyme, Q311-02) to perform qRT-PCR on a LightCycler 96 Real-Time PCR System (Roche). Rice *OsACTIN1* was used as the internal reference. All the primers used are shown in Table S2.

### ChIP assay

The ChIP assay was conducted as reported previously (Wu *et al*., 2021). Young panicles (0.5-3 cm) of the *35S:OsNLP3-FLAG* and *OsACTIN1:OsNLP4-GFP* transgenic plants grown in soil in a greenhouse were used for the ChIP assay. Specific antibodies anti-GFP (Abmart, M20004H) and anti-Flag (Abmart, M20008H) were used to precleared supernatants. Magnetic protein A/G beads (Bimake, B23202) were used to isolate antibody protein complexes. The enrichment of DNA fragments was quantified by qPCR using specific primers (Table S2). Enriched values were normalized to the level of input DNA. Wild type was used as a negative control.

### Electrophoretic mobility shift assay (EMSA)

EMSAs were performed as described previously (Yu et al., 2021). The coding regions of *OsNLP3*, *OsNLP4* and *OsRFL* were cloned into the reconstructive pET30a vector, containing an N-terminal MBP tag, His tag, and a Tobacco Etch Virus (TEV) protease site between the MBP tag and target protein. MBP-OsNLP3, MBP-OsNLP4, and MBP-OsRFL fusion proteins were expressed in the *Escherichia coli* Rosseta strain (DE3). We purified recombinant proteins and MBP protein by a Hisprep IMAC column (GE Healthcare). *Cis*-fragments of target gene promoters (34-bp free probes of *OsRFL* containing the NRE motif and 23-bp free probes of *OsMOC1* containing the WNNN (CCANTG) G/TNNNW *cis*-element) were synthesized, and biotin was labeled at the DNA ends. Unlabeled fragments of biotin with the same sequence were used as competitors. EMSAs were performed according to the LightShift^TM^ EMSA Optimization and Control Kit (Thermo Fisher Scientific, Waltham, MA, USA). MBP- OsNLP3- and MBP-OsNLP4-related reactions were loaded onto a 2% agarose gel in 0.5% TBE buffer for electrophoresis. MBP-OsRFL-related reactions were loaded onto a 6% native polyacrylamide gel in 0.5% TBE buffer for electrophoresis. The results were detected using a CCD camera system (Image Quant LAS 4000).

### Protoplast transfection and dual luciferase assay

The stems of rice grown in soil for 2-3 weeks were used to prepare protoplasts. In protoplast transient expression experiments, plasmids were transfected into protoplasts as described previously (Yu *et al*., 2021). The coding regions of *OsNLP3*, *OsNLP4* and *OsRFL* were cloned into pAN580 to generate effectors, respectively.

Approximately 3000 bp upstream of the start codons of *OsRFL* and *OsMOC1* were divided into three fragments on average, and each fragment was cloned into pGreenII 0800 to generate reporters. Firefly luciferase (LUC) activity and Renilla luciferase (REN) activity were determined 12-18 h after transfection using a dual luciferase reporting system (Vazyme, DL101-01). The LUC and REN activity ratios were determined at least three times.

### Statistical analysis

Statistical analysis was conducted by SPSS via one-way ANOVA with Tukey’s test or Student’s *t* test. Data are means ± SD. Different letters indicate a significant difference by Tukey’s test (*P* < 0.05). Asterisks indicate significant differences by Student’s *t* test (\**P* < 0.05, *\*\*P* < 0.01*, ***P* < 0.001).

## Data availability

The data supporting the findings of this study are available within this paper and its Supplementary information files. Sequence data of this paper can be found in the Rice Genome Annotation Project (http://rice.plantbiology.msu.edu/) under the following accession numbers: *OsNLP3 (LOC_Os01g13540)*, *OsNLP4 (LOC_Os09g37710)*, *OsRFL (LOC_Os04g51000)*, *OsMOC1 (LOC_Os06g40780)*, *OsMOC3 (LOC_Os04g56780)*, *OsLAX1 (LOC_Os01g61480)*, and *OsLAX2 (LOC_Os04g32510)*. The datasets and materials used during the current study are available from the corresponding author on reasonable request.

## Author Contributions

C.B.X. and J.W. designed the experiments. J.W., L.Q.S., Y.S., G.Y.W., J.X.W. and Y.B. performed the experiments and data analyses. J.W., Z.S.Z., J.Q.X., and Z.Y.Z. performed field trials and data analyses. L.Q.S. wrote the manuscript. C.B.X. and J.W. revised the manuscript. C.B.X. supervised the project.

## Acknowledgments

We thank Dr. Chengcai Chu (Institute of Genetics and Developmental Biology, Chinese Academy of Sciences) for the *35S:OsNLP3*-*FLAG* transgenic rice, Dr. Chuanzao Mao (College of Life Sciences, Zhejiang University), Dr. Shigui Li (State Key Laboratory of Hybrid Rice, Sichuan Agricultural University), Dr. Peng Qin (State Key Laboratory of Hybrid Rice, Sichuan Agricultural University) and Shimei Wang (Rice Research Institute, Anhui Academy of Agricultural Science) for their assistance with field trials, Dr. Qin Wang (Biotechnology Center, Anhui Agricultural University) for critical-point dryer support in the SEM assay, Dr. Chunming Wang (State Key Laboratory of Crop Genetics and Germplasm Enhancement, Nanjing Agricultural University) for vector pAN580, and Zhihao Xu (Division of Life Sciences and Medicine, University of Science and Technology of China) for assistance in protein purification.

## Supplementary figures and tables

**Figure S1.**
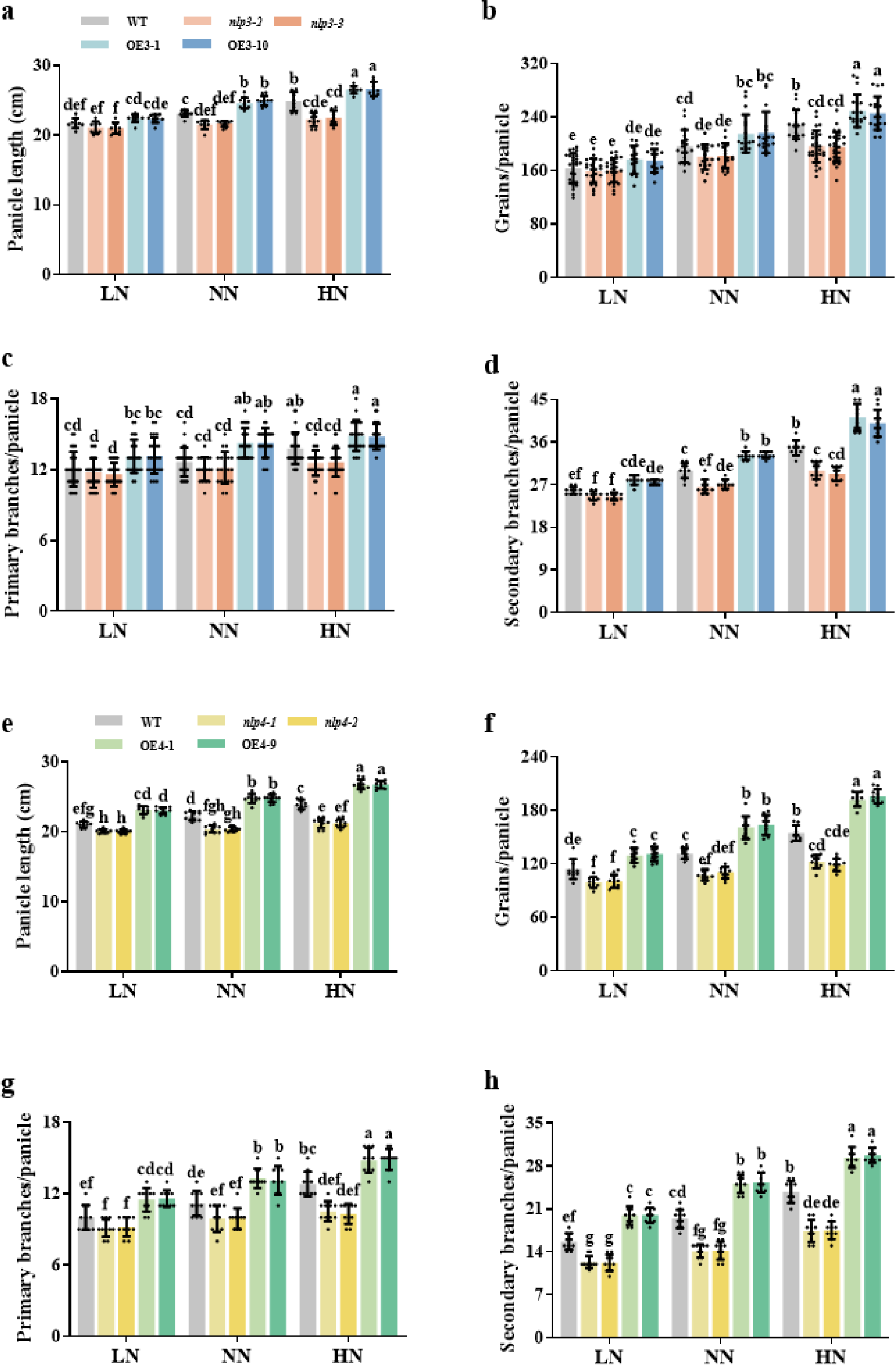
Panicle architecture-related traits of different *OsNLP3*/*4* genotypes under different nitrogen conditions. (**a-d**) Panicle architecture-related traits of the WT and different *OsNLP3* genotypes under low nitrogen (LN), normal nitrogen (NN), and high nitrogen (HN) conditions in Chengdu (April 2019 to September 2019), including panicle length (a), number of grains per panicle (b), number of primary branches per panicle (c), and number of secondary branches per panicle (d). Values are means ± SD (n ≥ 10). Different letters indicate significant differences by one-way ANOVA with Tukey’s test (*P* < 0.05). (**e-h**) Panicle architecture-related traits of the WT and different *OsNLP4* genotypes under LN, NN and HN conditions in Changxing (May 2019 to October 2019), including panicle length (e), number of grains per panicle (f), number of primary branches per panicle (g), and number of secondary branches per panicle (h). Values are means ± SD (n = 10). Different letters indicate significant differences by one-way ANOVA with Tukey’s test (*P* < 0.05).

**Figure S2.**
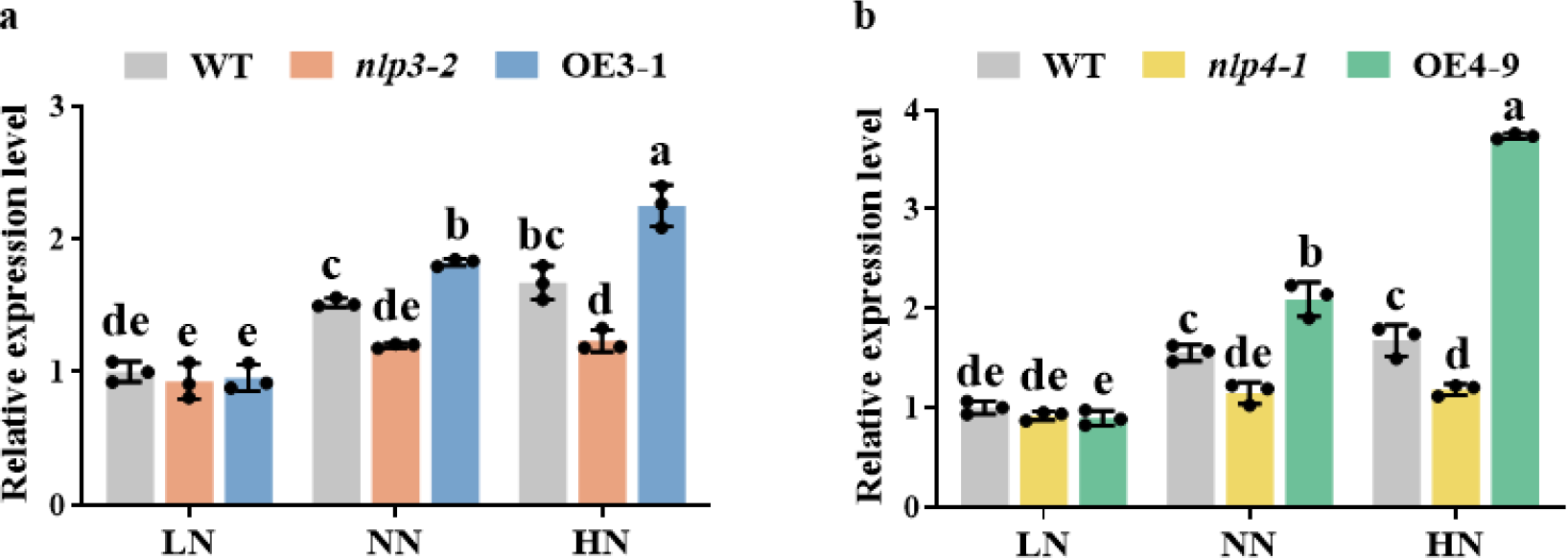
Relative expression levels of *OsRFL* in different *OsNLP3*/*4* genotypes under different nitrogen concentrations. Relative expression of *OsRFL* in different *OsNLP3* genotypes (a) and *OsNLP4* genotypes (b) grown in vermiculite pots under 0.5 mM nitrate (LN), 1 mM nitrate (NN), and 5 mM nitrate (HN) conditions. The expression levels are normalized to *OsACTIN1* at the branch meristem formation stage (0.5 cm young panicle). Values are means ± SD relative to WT under the LN conditions, which is set to 1 (n = 3). Different letters indicate significant differences by one-way ANOVA with Tukey’s test (*P* < 0.05).

**Figure S3.**
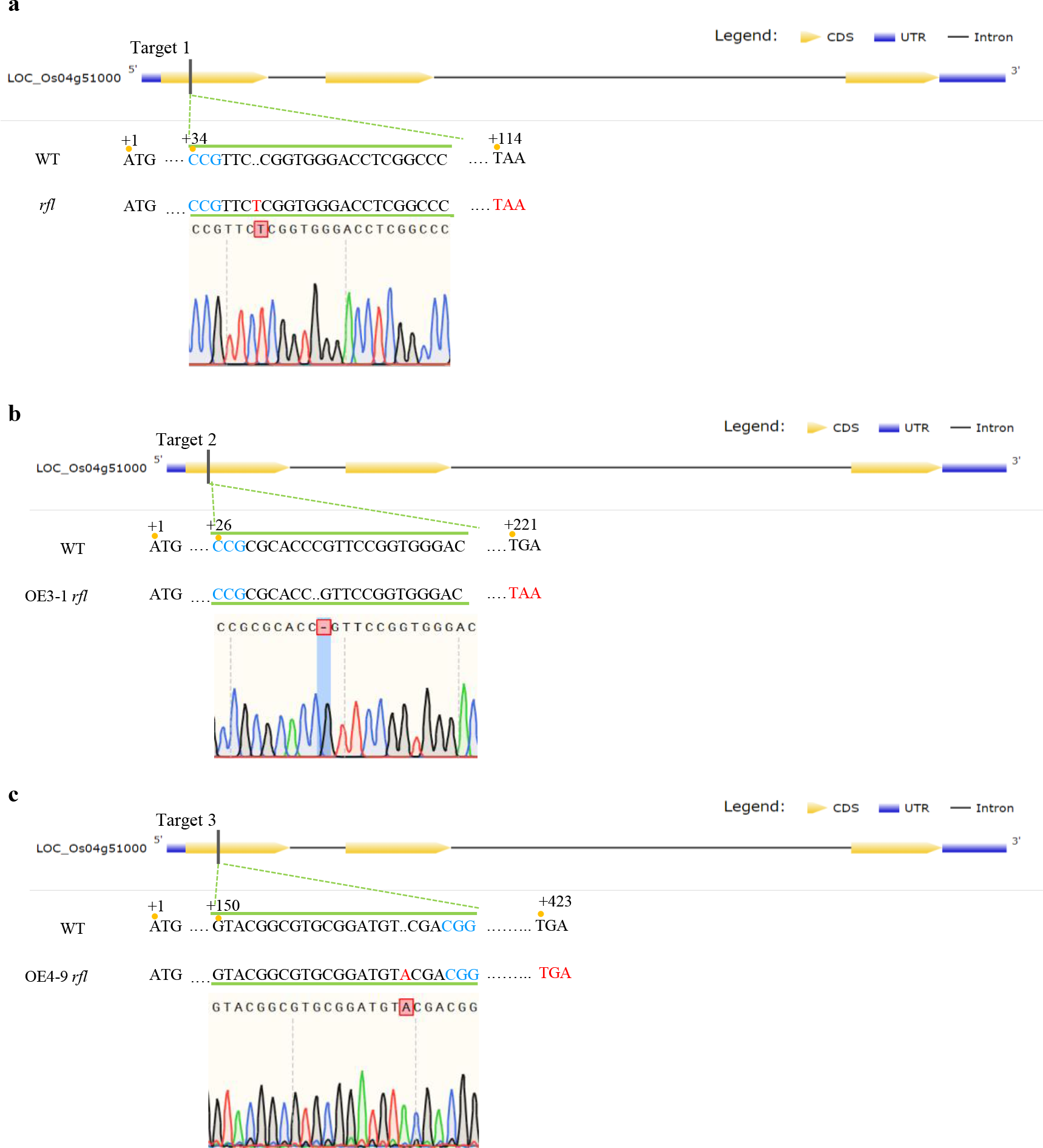
Verification of the CRISPR/Cas9-edited mutations of *OsRFL* in the *rfl*, OE3-1 *rfl*, and OE4-9 *rfl* mutants. Sequencing results of the edited *OsRFL* locus in *rfl* (a), OE3-1 *rfl* (b), and OE4-9 *rfl* (c) by CRISPR/Cas9. The *rfl* mutant has a T insertion at target 1, resulting in premature termination at exon 1. The OE3-1 *rfl* has a C deletion at target 2, resulting in premature termination at exon 1. The OE4-9 *rfl* has an A insertion at target 3, causing a premature termination in exon 2.

**Figure S4.**
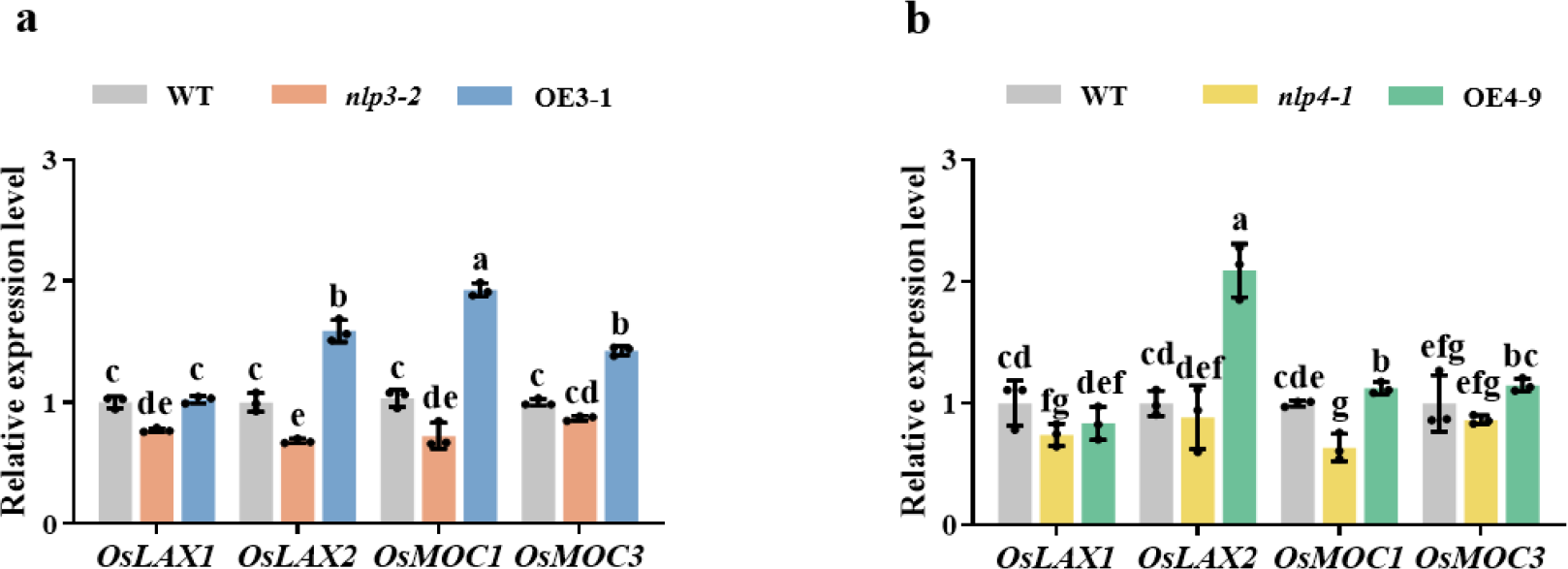
Relative expression levels of IM maintenance and specification marker genes in different *OsNLP3*/*4* genotypes. Relative expression of *OsLAX1*, *OsLAX2*, *OsMOC1* and *OsMOC3* in different *OsNLP3* genotypes (a) and *OsNLP4* genotypes (b), normalized to *OsACTIN1* at the branch meristem formation stage (0.5 cm young panicle). Values are means ± SD relative to the WT which is set to 1 (n = 3). Different letters indicate significant differences by one-way ANOVA with Tukey’s test (*P* < 0.05).

**Table S1.** Comparison of panicle architecture-related traits of different *OsNLP3*/*4* genotypes under different nitrogen conditions.

| Trait | Panicle length |  |  | Grain number/panicle |  |  | Primary branch/panicle |  |  | Secondary branch/panicle |  |  |
| --- | --- | --- | --- | --- | --- | --- | --- | --- | --- | --- | --- | --- |
|  | LN | NN | HN | LN | NN | HN | LN | NN | HN | LN | NN | HN |
| *nlp3* | -10.35% | -11.42% | -12.02% | -18.28% | -21.27% | -25.53% | -15.63% | -17.16% | -20.75% | -14.78% | -18.84% | -21.91% |
| **OE3** | +12.43% | +13.78% | +14.25% | +13.12% | +18.79% | +26.18% | +14.58% | +18.93% | +19.42% | +5.17% | +10.92% | +15.36% |
| *nlp4* | -9.57% | -9.40% | -10.53% | -16.68% | -21.57% | -22.99% | -4.01% | -7.10% | -12.19% | -18.93% | -22.08% | -28.04% |
| **OE4** | +7.38% | +12.05% | +13.27% | +21.79% | +26.39% | +40.00% | +22.99% | +27.70% | +40.62% | +27.02% | +37.92% | +45.63% |

The average values of the panicle architecture-related traits for different *OsNLP3/4* genotypes compared to the WT were calculated. *nlp3* represents the *OsNLP3* knockout mutants (*nlp3-2* and *nlp3-3*), OE3 represents the *OsNLP3* overexpression lines (OE3-1 and OE3-10), *nlp4* represents the *OsNLP4* knockout mutants (*nlp4-1* and *nlp4-2*), and OE4 represents the *OsNLP4* overexpression lines (OE4-1 and OE4-9). LN represents low nitrogen conditions (80 kg urea per ha for *OsNLP3* genotypes, 90 kg urea per ha for *OsNLP4* genotypes), NN represents normal nitrogen conditions (160 kg urea per ha for *OsNLP3* genotypes, 180 kg urea per ha for *OsNLP4* genotypes), and HN represents high nitrogen conditions (320 kg urea per ha for *OsNLP3* genotypes, 360 kg urea per ha for *OsNLP4* genotypes).

**Table S2.**
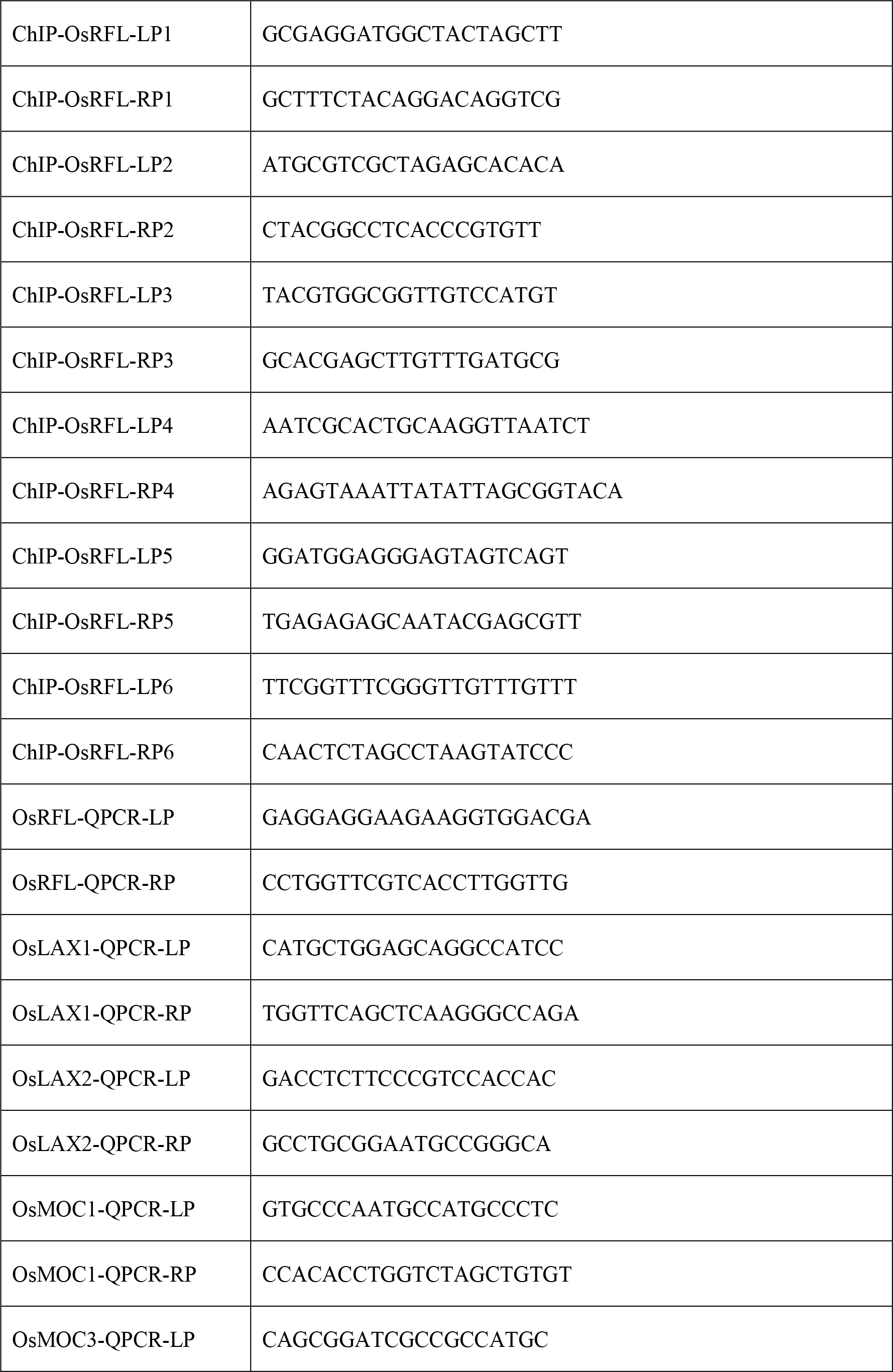
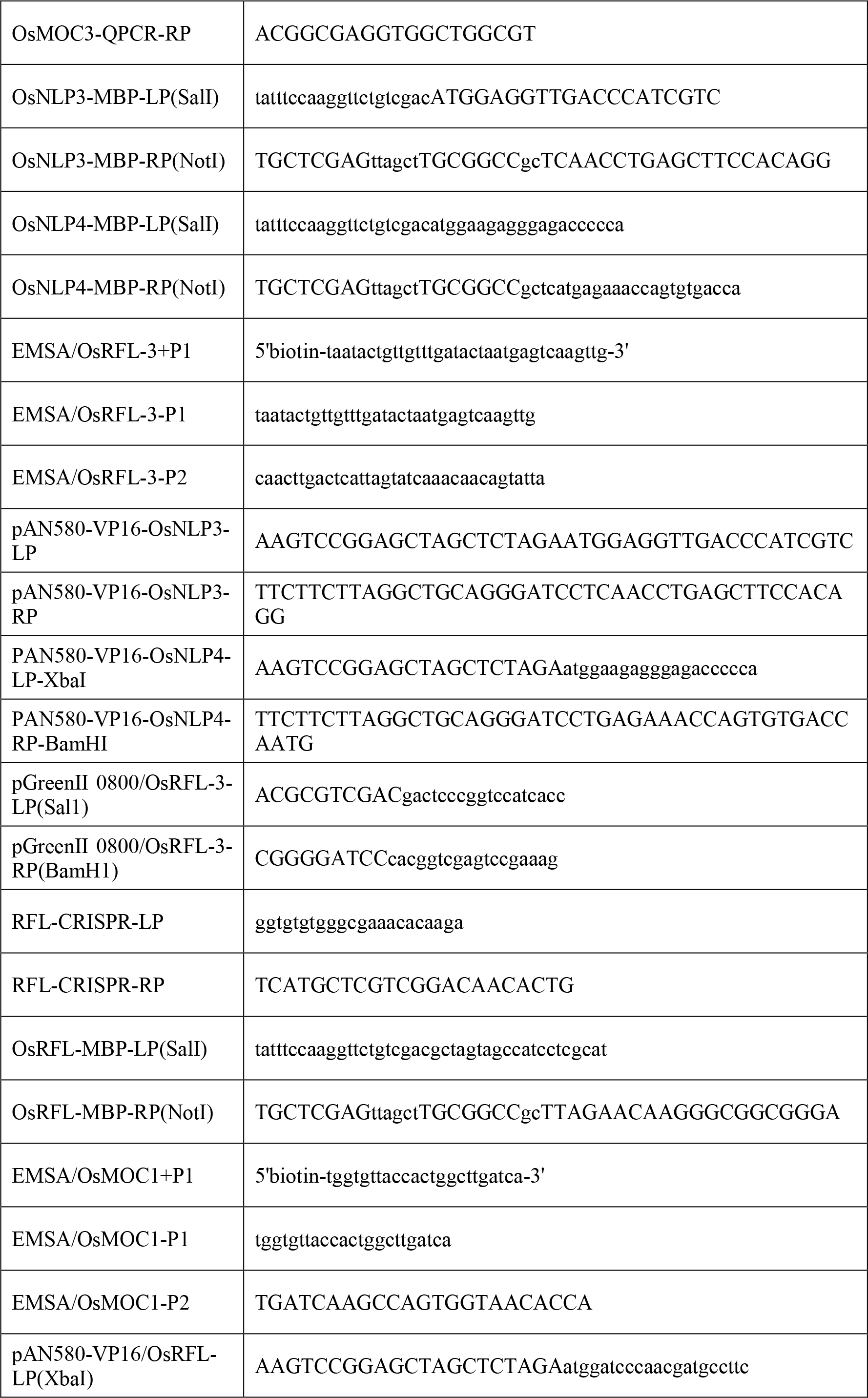
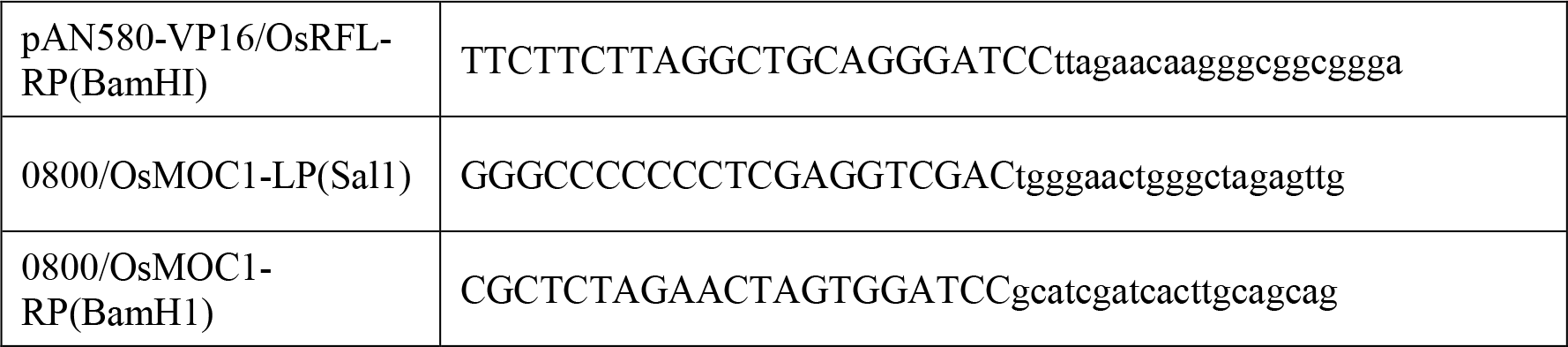
List of primers used in this study.

